# Identifying transcription factors driving cell differentiation

**DOI:** 10.1101/2025.01.20.633856

**Authors:** Paul Martini, Anne Hartebrodt, Dietmar Zehn, David B. Blumenthal

## Abstract

Cellular differentiation is a fundamental biological process at the core of development and growth in multicellular organisms. Understanding it at the level of transcriptomic regulation opens the possibility for novel biological insights, for steering cell development for therapeutic purposes, and for developing targeted therapies for diseases. While many methods exist that infer developmental trajectories from single-cell RNA sequencing data, only few can effectively determine which biological mechanisms drive differentiation along such trajectories. To close this gap, we developed SwitchTFI (**switch t**ranscription **f**actor **i**dentification), a method to identify differentiation-driving regulatory mechanisms and the transcription factors (TFs) that play a key role in them. SwitchTFI infers a cell state transition-specific gene regulatory network (GRN) from a user-provided baseline GRN via decision stump learning and permutation-based significance tests. Key TFs are then extracted by ranking them according to their centralities in the transition GRN. We show that SwitchTFI can identify TFs known to be involved in pancreatic endocrinogenesis and in erythrocyte differentiation and that it outperforms competitor methods with respect to the functional coherence of the predicted driver TFs. SwitchTFI is available as a Python package at https://github.com/bionetslab/SwitchTFI.

## 1 Introduction

The process of cell differentiation is mirrored in the transcriptional landscape [Carninci et al., 2005], where cells navigate by committing to lineage trajectories, leading to their distinct cellular identity and function. Single-cell RNA sequencing (scRNA-seq) provides a detailed view at the transcriptomic profile of individual cells at different development stages and is thus a great tool for elucidating cell differentiation. However, scRNA-seq data is usually captured at a single time point, making the reconstructing of the developmental trajectories an important problem in data-driven systems biology. While numerous methods for computational trajectory inference exist [Trapnell et al., 2014, Setty et al., 2016, Street et al., 2018, Setty et al., 2019, Lange et al., 2022], relatively little work has been done in developing methods that determine the biological mechanisms and key transcription factors (TFs) that drive cell differentiation along an identified trajectory. Notable exceptions are the tools CellRank [Lange et al., 2022] (an improved version of Palantir [Setty et al., 2019]), spliceJAC [Bocci et al., 2022], and DrivAER [Simon et al., 2020]. However, all of these tools are subject to important limitations, as summarized in the following paragraphs (see Section 1 of the supplement for further details).

Palantir identifies branches of the developmental trajectory by analyzing the properties of a Markov process on a nearest-neighbour graph embedding of the scRNA-seq data. This yields a probability distribution over the identified branches for each cell. The Pearson correlation between the expression of individual genes and branch probabilities is used to identify driver genes. CellRank improves upon Palantir by integrating RNA velocity [La Manno et al., 2018, Bergen et al., 2020] for inferring the directionality of the edges in the graph embedding. Both methods suffer from the risk of encountering spurious correlations, since no functionality is provided to test the obtained correlations for statistical significance.

The spliceJAC method assumes a scRNA-seq count matrix with spliced and unspliced counts and a cell annotation vector with the respective cell states as an input to infer cell state-specific gene regulatory networks (GRNs), using a dynamical system model. Genes that are critical to the transition between cell states are found by analyzing the eigenspace of the Jacobian matrix associated with the differential equations of the starting cell state. By construction, the number of considered genes must be limited to the number of cells within a given cell state, such that an extensive preselection is required. In the tutorial published with the spliceJAC Python package, only 50 out of 27998 possible genes are used for GRN inference and transition driver gene identification. Notably, spliceJAC also provides a transition-specific GRN. However, it is constructed *post ho*c, using previously inferred marker and driver genes as vertices and the interactions between them as edges. Since no transition-specific information is utilized for predicting these interactions, their relevance to the cell state transition is unclear. Further, it is unclear how a sensible cutoff for the vertices and edges to be included in the GRN should be defined.

Given a scRNA-seq count matrix, a family of gene sets, and a phenotype vector of interest, DrivAER combines a deep count autoencoder network [Eraslan et al., 2019] for denoising and imputation of scRNA-seq data with a random forest model to compute relevance scores for the individual gene sets. TFs that drive differentiation can be found by defining the family of gene sets as a collection of sets of target genes of known TFs and using cell state annotations as the phenotype vector. Thus, mechanistic information is directly utilized for driver TF identification. However, since DrivAER works on the scale of TFs and their corresponding sets of target genes, it cannot recover which individual links between TFs and target genes are relevant.

To overcome the limitations of existing methods, we present **SwitchTFI** (**switch t**ranscription **f**actor **i**dentification) (see Table 1 for a high-level feature comparison against existing methods). SwitchTFI leverages the combined information of a scRNA-seq dataset annotated with progenitor and offspring cell states and a baseline GRN from an appropriate biological context to compute a transition GRN as an interpretable model of the regulatory mechanisms that drive cell state transitions. To infer the transition GRN from the baseline GRN, we define cell state transition relevance weights for all edges in the baseline GRN and then use a variant of the Westfall-Young permutation method [Westfall and Stanley Young, 1993] for family-wise error rate (FWER) correction to only keep those edges where these weights are significant. Subsequently, we use node centrality measures to identify key TFs within the transition GRN.

**Table 1.** Features of SwitchTFI and other methods for identifying differentiation driver genes.

| Method | CellRank | spliceJAC | DrivAER | SwitchTFI |
| --- | --- | --- | --- | --- |
| Publication | Lange et al. [2022] | Bocci et al. [2022] | Simon et al. [2020] | This article |
| Input | scRNA-seq count matrix with spliced and unspliced counts or with pseudotime annotations | scRNA-seq count matrix with spliced and unspliced counts and with cell state annotations | scRNA-seq count matrix with cell state annotations and baseline GRN | scRNAseq count matrix with cell state annotations and baseline GRN |
| Output | Ranked genes | Ranked genes and transition GRN | Ranked TFs | Ranked TFs and transition GRN |
| Unit of analysis | Genes | Genes | TFs | Edges between TFs and target genes |
| Inbuilt statistical validation | None | None | None | Edge-level $P$-values |

## 2 Results

### 2.1 Inferring transition GRNs with SwitchTFI

An overview of the SwitchTFI method is provided in Figure 1 (see Methods for details). As input, SwitchTFI requires scRNA-seq data from cells of annotated consecutive cell states (hereafter referred to as “progenitor” and “offspring” cells) and a baseline GRN containing the edges from TFs to target genes. For this study, we used Scenic [Aibar et al., 2017] to infer the baseline GRN from the scRNA-seq data. Alternatively, expert-curated TF-target gene links obtained from databases such as CollecTRI [Müller-Dott et al., 2023] could be used as baseline GRN. For each edge *e* of the baseline GRN, we predict the target gene expression from the expression of the TF, using a regression tree of depth one. The regression tree’s decision boundary induces a partition of the cells. We compare this partition to the progenitor-offspring partition provided by the cell state annotations and obtain an edge weight *w*_*e*_ *∈* [0, 1] that quantifies similarity between the two partitions. SwitchTFI’s key idea is that the two partitions are similar if the TF-target gene link is relevant to the cell state transition (i.e., if the TF drives differentiation). In this case, the cell state annotations induce a distinct clustering of the cells in TF-target gene expression space, leading to a large edge weight *w*_*e*_.

**Figure 1.**
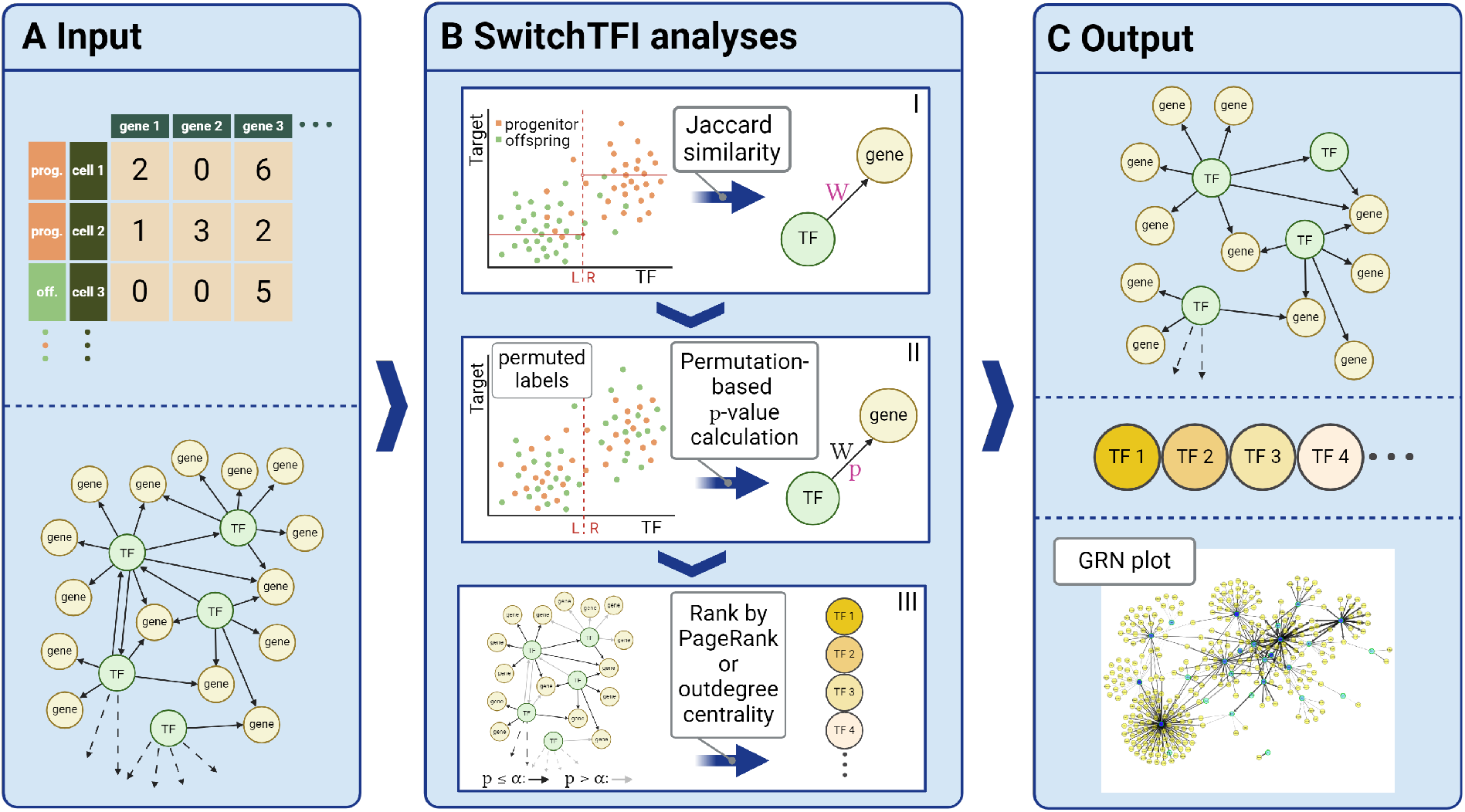
Overview of SwitchTFI. (A) Input: a scRNA-seq count matrix with progenitor/offspring annotations and GRN from a fitting biological context (possibly inferred from the scRNA-seq data). (B) Workflow: (I) weight fitting, (II) empirical *P*-value calculation, and (III) pruning of the GRN and ranking of TFs according to centrality in the transition GRN. (C) Output: transition GRN, ranked list of driver TFs, and plot of the transition GRN.

By adopting this edgewise approach instead of analyzing single genes, SwitchTFI focuses on the smallest regulatory unit that still carries mechanistic information, i.e., TF-target gene links. To capture the regulatory mechanisms that drive differentiation on the scale of the GRN, SwitchTFI prunes edges from the baseline GRN, retaining only the ones most relevant to the cell state transition in a transition GRN. For this, we permute the cell state annotations, compute edge weights 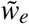 for the permuted cell states, and then compute adjusted Westfall-Young *P*-values *P*_*e*_ to assess if *w*_*e*_ is significantly larger than expected by chance. By retaining only edges with *P*_*e*_ *≤ α*, we obtain a transition GRN that is specific to the respective cell state transition with controlled FWER *≤ α*.

The transition GRN is a highly interpretable output format that is well suited for further computational and biological analyses. It can be visualized with the convenient plotting functionality provided by the Python implementation of SwitchTFI. Similar to existing methods, SwitchTFI also returns a ranked list of putative key driver TFs. SwitchTFI provides two methods to compute this ranked list: PageRank centrality [Brin and Page, 1998], which is widely used for gene ranking in computational systems biology [Ding et al., 2021, Xu et al., 2024], and weighted outdegree centrality as a conceptually simpler and more interpretable alternative. For both options, the TF ranking is hence mechanistically informed, as its computation is based on the topology of the transition GRN. Together with the transition GRN, the key TFs can be interpreted within their broader regulatory context.

For the sake of a concise presentation, we used PageRank centrality for our in-depth analyses reported in Sections 2.2 and 2.3. For the comparison against existing methods in Section 2.4, we tested both PageRank and weighted outdegree centrality. The choice of the FWER threshold *α* mainly influences the size of the transition GRN, while the ranking of the TFs remains largely unchanged (see Section 3 of the supplement). For all results reported in this article, we chose *α* = 0.05, which results in a concise and well-interpretable output.

### 2.2 Driver TFs identified by SwitchTFI and corresponding target genes show characteristic dynamics over pseudotime

To validate SwitchTFI and examine the validity of its underlying assumptions, we ran it on a scRNA-seq dataset of murine pancreatic tissue from embryonic day 15.5 from Bastidas-Ponce et al. [2019]. Moreover, we used the erythrocyte differentiation dataset by Paul et al. [2015]. Pancreatic endocrinogenesis and haematopoiesis are well-studied processes that are often used as templates for developing and testing computational methods that aim at elucidating cellular differentiation [Bergen et al., 2020, Lange et al., 2022, Bocci et al., 2022, Wolf et al., 2019]. For details on the datasets and their availability, see Section 4.5.

Figure 2 provides a visual overview of SwitchTFI’s results on the *β*-cell development dataset. For highly weighted edges, the cell state annotations have a pronounced cluster structure that corresponds well to the partition induced by the regression tree (Figure 2A). The converse can be observed for lowly weighted edges (Figure 2B). Further, we looked at the shape of gene expression trends in pseudotime. Genes with strongly decreasing or increasing expression in pseudotime are generally adjacent to edges with a high weight (Figure 2A, C, D). Nearly constant expression levels along the pseudotemporal trajectory are associated with low weights (Figure 2B, E, F). Genes with rapid expression changes during differentiation are of interest for further investigation, since such changes can be linked to governing differentiation and changes in cell function. For the top 10 putative driver TFs a rapid, switch-like increase or decrease in expression can be observed (Figure 2G). This behavior carries over to the targets of the top TFs. For instance, Pdx1 is a TF that exhibits a marked increase in expression over pseudotime. The same can be observed for its targets (Figure 2H), indicating the importance of the Pdx1 regulatory complex for developed *β*-cells. Similarly, the diminishing expression of Pax4 over pseudotime coincides with the downregulation of its targets (Figure 2I). Thus, we can associate Pax4 with inducing and regulating differentiation towards the *β*-cell fate but declining importance to mature *β*-cells. Overall, these results show that SwitchTFI’s underlying methodological assumption that highly weighted edges point towards regulatory links being involved in cell state transition is reasonable.

**Figure 2.**
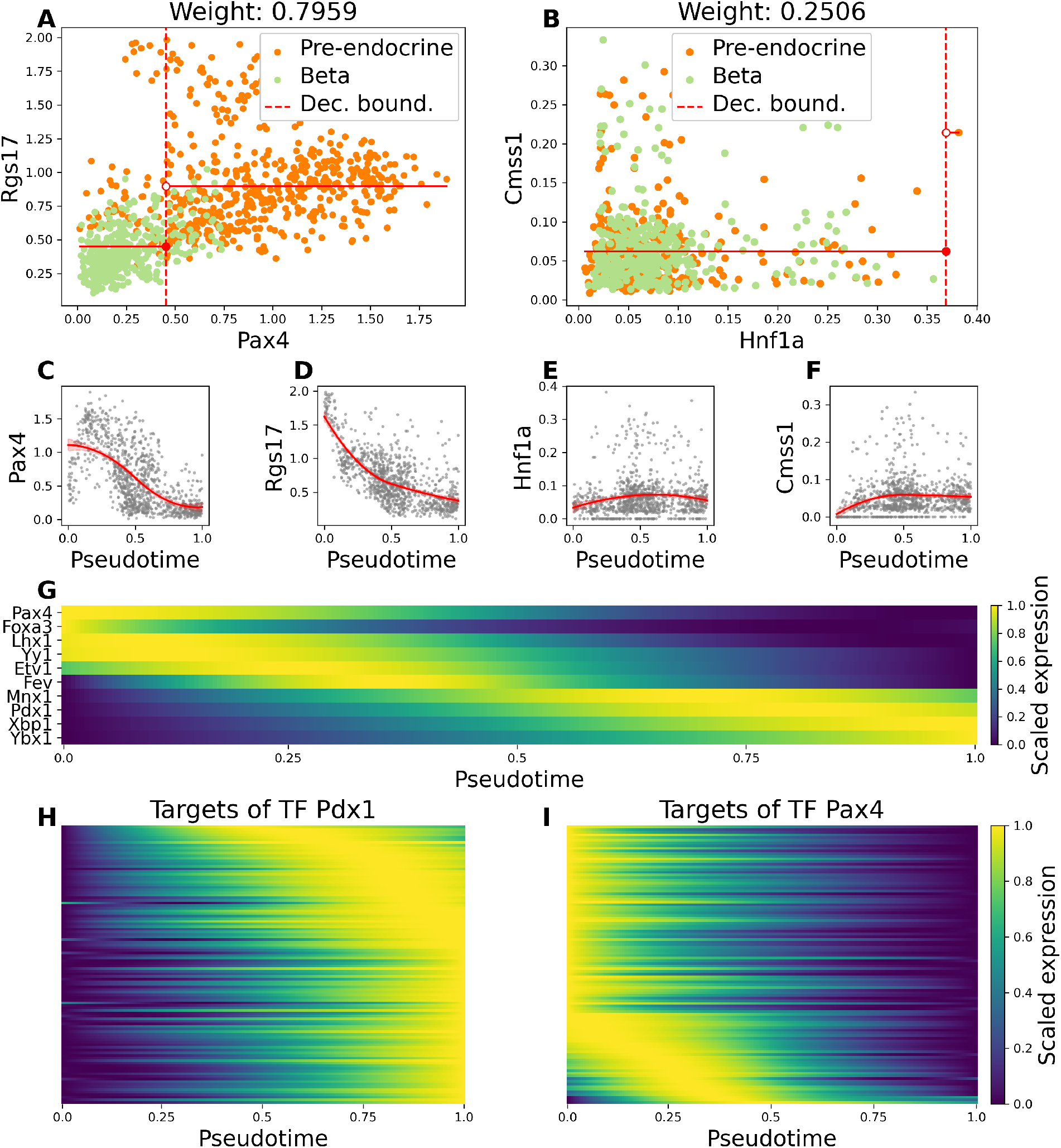
Gene expression of key TFs and target genes uncovered by SwitchTFI on the mouse *β*-cell development data. (A, B) Scatter plots of TF expression (*x*-axis) and target gene expression (*y*-axis) of the cells in the scRNA-seq dataset. (A) corresponds to a highly and (B) to a lowly weighted edge. The regression stump fit during the weight fitting step is plotted in red. (C – F) Gene expression trends over pseudotime for the individual TFs and target genes of the edges shown in (A) and (B). (G) Expression trends over pseudotime for the top-10 TFs predicted to be relevant to the cell state transition. TFs have been sorted according to their trend’s peak in pseudotime. (H, I) Expression trends over pseudotime for the targets regulated by putative driver TFs Pdx1 and Pax4.

In addition to the qualitative validation shown in Figure 2, we quantitatively examined SwitchTFI’s output with respect to the fraction of genes with biologically interesting, rapidly changing expression trends. To quantify a rapid change in gene expression, we tested for differential expression along a previously computed pseudotemporal trajectory (ptDE). A ptDE *q*-value was computed for each gene of the baseline GRN using switchde [Campbell and Yau, 2017]. By pairwise combining the *q*-values of TFs and targets using Fisher’s method, we obtained edgewise ptDE *q*-values. Figure 3A and D visualize the edge weights and the combined log-transformed *q*-values for all edges of the unpruned baseline GRN computed by Scenic. Edges below the *q*-value threshold are deemed insignificant with respect to ptDE. Edges above the weight threshold have an adjusted permutation-based *P*-value below 0.05 and are thus retained in the transition GRN. Notably, all such edges have a significant combined *q*-value (*≤* 0.01), and we observe a clear correlation between large weights and small combined *q*-values (Pearson correlation coefficient between non log-transformed *q*-values and weights for *β*-cell transition: *−*0.258, erythrocytes: *−*0.131).

**Figure 3.**
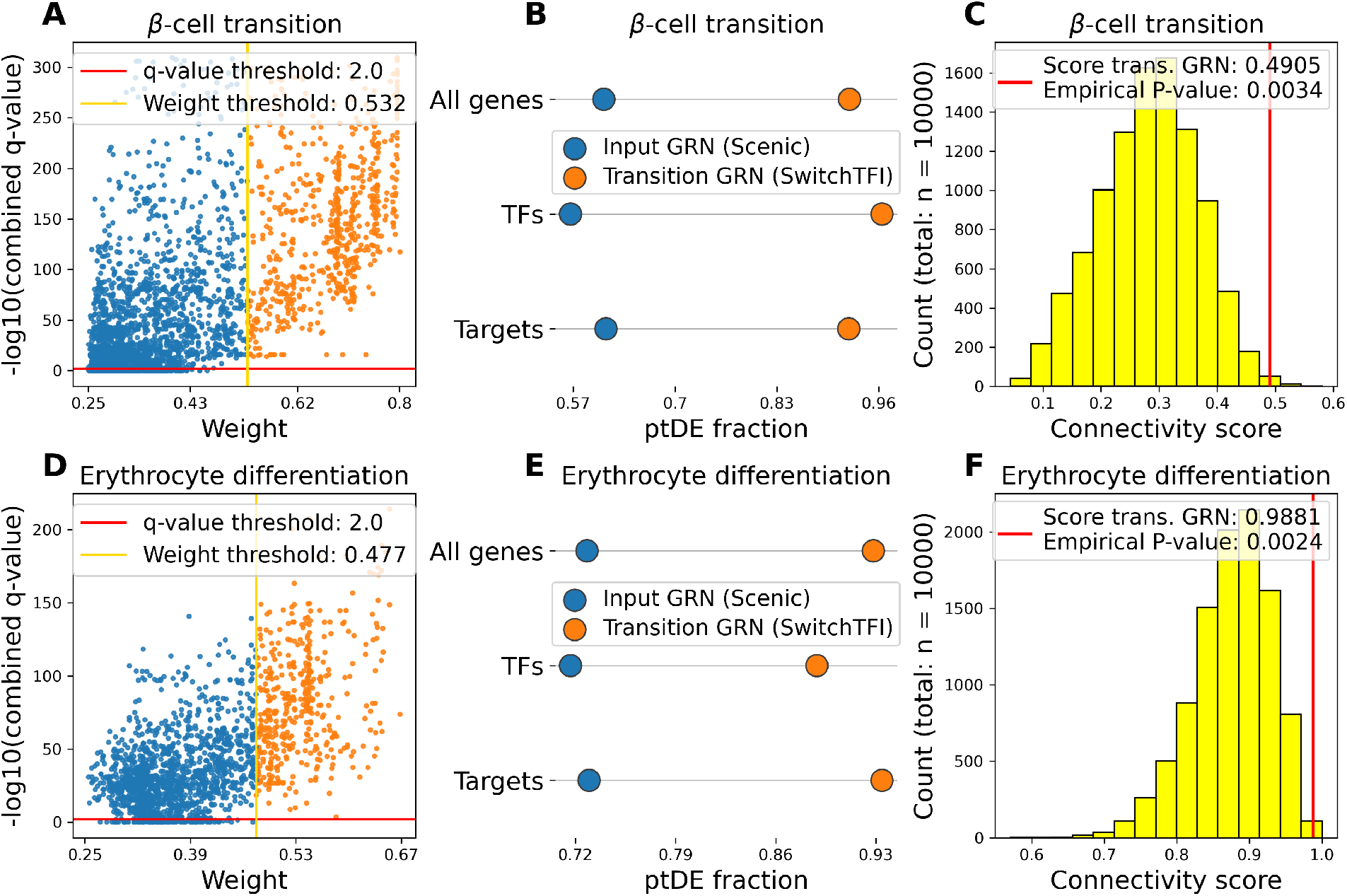
Quantitative inspection of SwitchTFI’s transition GRNs for the *β*-cell transition and the erythorcyte differentiation data (see Supplementary Figure 1 for corresponding plots for the *α*-cell transition dataset). (A, D) Scatter plot of edge weights vs. combined ptDE *q*-values of the unpruned input GRN computed by Scenic. Combined *q*-values are *−* log_10_(*·*)-transformed. (B, E) Fraction of ptDE genes, TFs and targets in the Scenic and SwitchTFI GRN. (C, F) Histogram of the connectivity scores of randomly sampled subnetworks of the Scenic GRN. The transition GRN’s score is visualized as a red line.

Figure 3A and D show that combined *q*-values are a relatively broad criterion for quantifying the relevance of an edge in the baseline GRN to the cell state transition, as only few edges do not have a significant combined *q*-value. Although the SwitchTFI algorithm does not rely on the combined *q*-values, we observe that SwitchTFI *de facto* makes a mechanistically informed selection among the edges with a significant ptDE combined *q*-value. The biological relevance of this selection is confirmed by the observation that the fraction of ptDE genes, TFs, and target genes in the transition GRN is substantially increased compared to the baseline GRN (Figure 3B, E).

Further, we compared the number and size of the connected components in SwitchTFI’s transition GRNs against randomly sampled subnetworks of the baseline GRN. As a score for the comparison, we used the sum over the squared ratios of connected component sizes to the total number of vertices in the subnetwork. This way, more tightly connected subnetworks in which many nodes are concentrated in few connected components receive a higher connectivity score (for details see Section 4.12). Figure 3C and Figure 3F show that the transition GRNs consist of significantly fewer and larger connected components than expected under the random background model, indicating that SwitchTFI’s transition GRNs indeed comprise biologically relevant internally connected regulatory units.

### 2.3 SwitchTFI recovers key transcription factors involved in pancreatic endocinogenesis

In Figure 4, the top 10 pre-endocrine *β*- and *α*-cell driver TFs as ranked by SwitchTFI are shown (Figure 4A), together with the expression trends of the *α*-cell drivers over pseudotime (Figure 4B, see Figure 2G above for the *β*-cell drivers’ expression trends over pseudotime). For both lists of driver TFs, we performed gene set enrichment analysis, using the Enrichr webtool [Chen et al., 2013, Kuleshov et al., 2016]. For the *β*-cell drivers, a significant enrichment can be observed for terms that relate to general regulation of transcription or more specifically regulation of *β*-cell development (Figure 4C, see Supplementary Figure 1 for enrichment results for *α*-cell drivers), indicating that key TFs retrieved by SwitchTFI are functionally relevant. Moreover, for several of the top-ranked TFs, there is strong literature support for their involvement in pancreatic endocrinogenesis:

- Pdx1 appears among the top-ranked genes for the *β*- and *α*-transition. This is unsurprising, since Pdx1 assumes a central role in governing endocrinogenesis [Dassaye et al., 2016, Oliver-Krasinski et al., 2009, Wang et al., 2001, Ebrahim et al., 2022] and its expression is essential to the development of the endocrine system [Dassaye et al., 2016]. Cellular reprogramming of *β*-cells to *α*-like cells by Pdx1-deletion was experimentally confirmed [Gao et al., 2014]. Our results fully confirm this antithetical relevance of Pdx1 to differentiating *β*- and *α*-cells (see contrary expression trends over pseudotime in Figure 2G and Figure 4B).
- The TF Xbp1 is highly ranked as a driver of the *β*-cell transition and shows an increasing expression over pseudotime (Figure 2G). Xbp1 has been linked to differentiation processes in many cell types, including *β*-cells. In particular, it was shown that Xbp1 has a key role in maintaining *β*-cell identity and repressing transdifferentiation into other pancreatic islet cells, such as *α*-cells [Lee et al., 2022].
- Arx is identified as key TF driving *α*-cell development and exhibits an increasing expression over pseudotime (Figure 4B). This is coherent with the finding that Arx expression is sufficient for maintaining *α*-cell identity in the developing and neonatal pancreas [Collombat et al., 2007, Wilcox et al., 2013].
- Pax4 appears among the top 10 drivers inferred for both *α*- and *β*-cell development and, for both trajectories, its expression peaks early in pseudotime (Figure 2G, Figure 4B). These findings are in line with experimental studies according to which Pax4 plays an important role in the regulation of pancreatic endocrine development [Collombat et al., 2003, Dassaye et al., 2016] but alone does not suffice to determine endocrine subtype fate [Greenwood et al., 2007].

**Figure 4.**
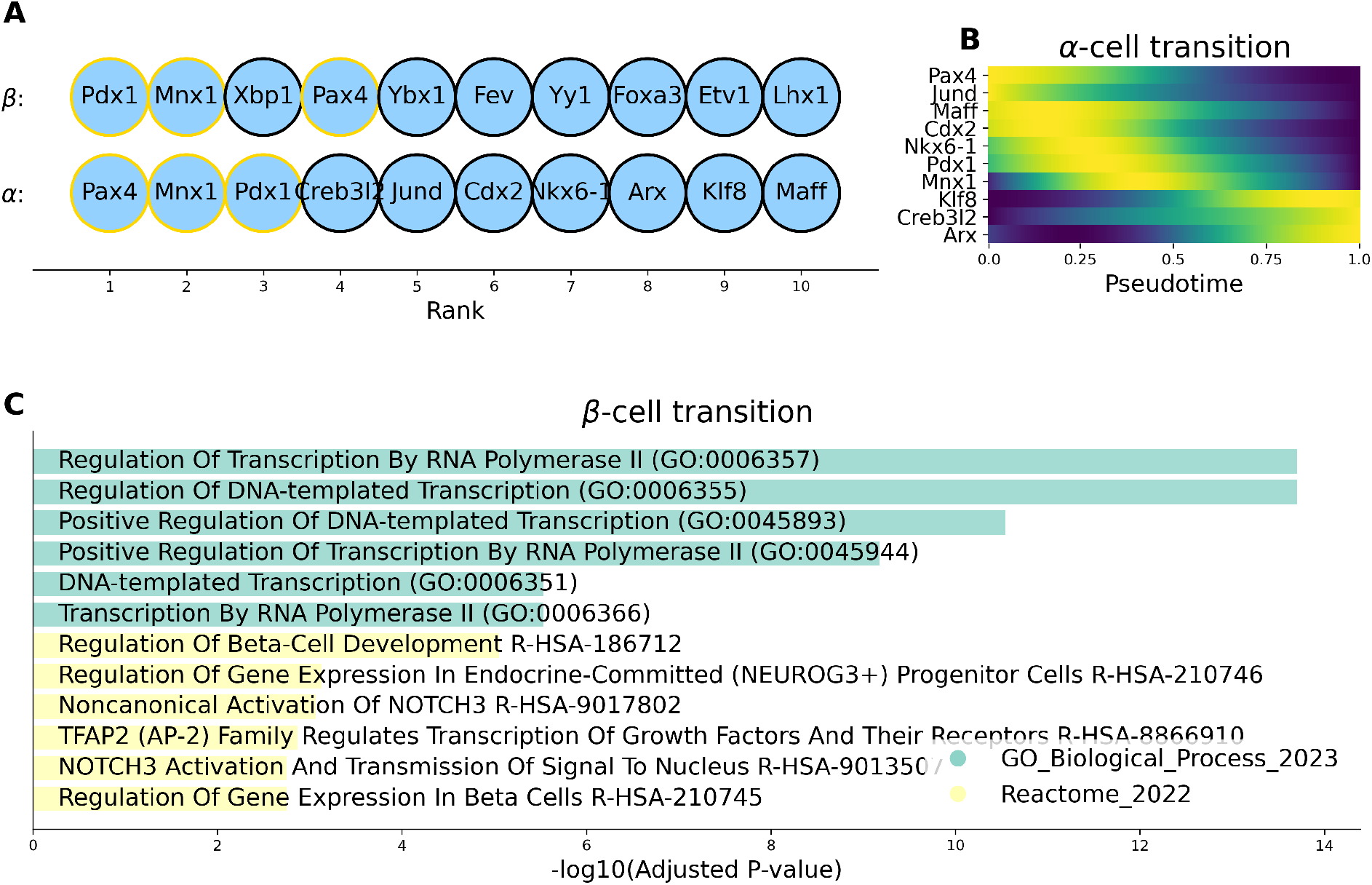
Qualitative validation of the top 10 pre-endocrine *β*-/*α*-cell transition driver TFs, as ranked by SwitchTFI (see Supplementary Figure 2 for corresponding results for the erythrocyte differentiation data). (A) Rankings of the putative driver TFs. Yellow outline marks TFs appearing in both rankings. (B) Expression trends in pseudotime of the top 10 TFs predicted to drive the transition to *α*-cells. TFs have been sorted according to their trends’ peaks in pseudotime. (C) Gene set enrichment results for the top 10 putative *β* driver TFs. The top 6 terms of each reference database are ranked according to their adjusted *P*-value as computed by Enrichr.

Beyond these TFs which are known to be involved in pancreatic endocrinogenesis, SwitchTFI predicts Ybx1 as the fifth most important TF for *β*-cell development (Figure 4A). Prior research has shown that Ybx1 is an important modulator of epithelial cell differentiation [Kwon et al., 2018]. Ybx1 is also implicated in various disease pathways, where it appears to modulate epitelial-menenchymal transition [Guo et al., 2022, Kwon and Kim, 2024, Bai et al., 2023], highlighting the potential implication of this gene in differentiation processes. Thus, we examined Ybx1’s top 20 predicted targets in the transition GRN (Figure 5A). Notably, two of these targets, Meg3 and Tmed10, have been linked to exocrine pancreatic differentiation in prior studies, especially in the context of diabetes mellitus [Li et al., 2022, Zhou et al., 2023, Tao et al., 2023]. Gene set enrichment analysis of the top 20 targets returned a collection of terms related to oxidative processes and energy metabolism (Figure 5B). Ox-idative stress due to reactive oxygen species (ROS) plays an important role in *β*-cell differentiation and maintenance [Wang and Wang, 2017]. ROS and mitochondrial deficiencies have also been linked to *β*-cell dysfunction and failure in type II diabetes (T2D) [Haythorne et al., 2019, Onikanni et al., 2023]. Therefore, one could hypothesize that Ybx1 is implicated in the differentiation or maintenance of *β*-cells via the modulation of genes controlling the oxidative environment.

**Figure 5.**
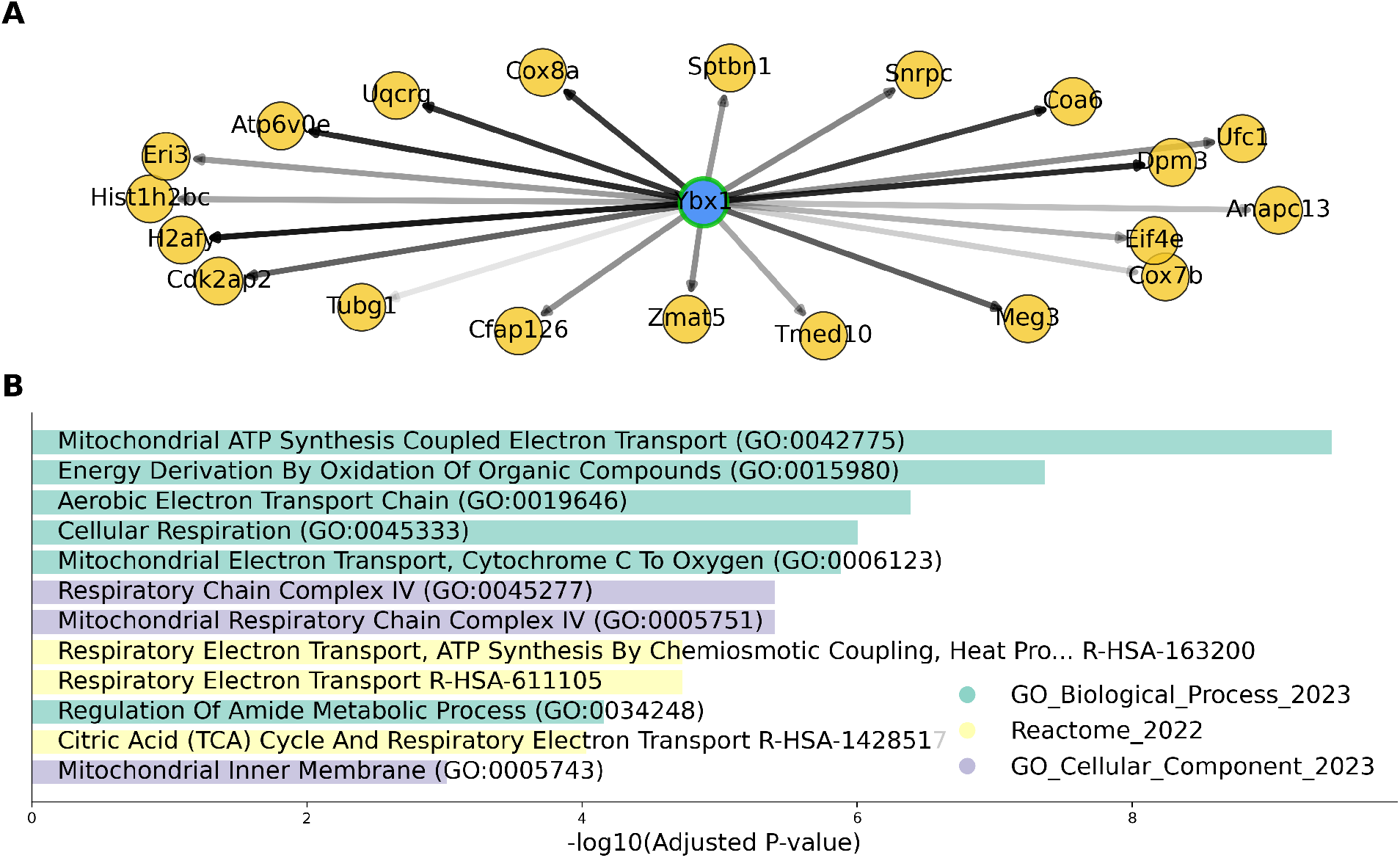
Exemplary hypothesis generation with SwitchTFI. (A) TF Ybx1 and its top 20 target genes, ranked according to SwitchTFI’s edge score computed from the edge weight and the empirical *P*-value (see Section 4.4). The shade of the edges is proportional to the scores, darker shades indicate higher relevance. The network plot was generated with SwitchTFI’s inbuilt plotting function `switchtfi.plotting.plot regulon()`, which supports out-of-the-box visualization of inferred regulons. (B) Gene set enrichment results for the top 20 targets of TF Ybx1. Terms are ranked according to their adjusted *P*-value as computed by Enrichr.

In sum, SwitchTFI is hence able to recover TFs that are biologically validated to drive differentiation towards known endocrine cell fates. Moreover, the example of the Ybx1 regulon showcases SwitchTFI’s potential for data-driven hypothesis generation in settings where the user has prior evidence on genes (Meg3 and Tmed10 in our example) involved in a cell state transition of interest and aims to identify candidate upstream regulators (here: Ybx1) and their action mechanisms to be validated in follow-up wet lab experiments.

### 2.4 Comparison of SwitchTFI to other methods

To compare the results of SwitchTFI to the previously discussed competitor methods, we ran each on the pre-endocrine *β*-cell transition data and on the erythrocyte differentiation data. For the corresponding results for the pre-endocrine *α*-cell transition data, we refer to Supplementary Figure 1. SwitchTFI was run in two modes: with TFs ranked according to PageRank and according to their weighted outdegree in the transition GRN. Since only spliceJAC has a GRN as part of its output, we resorted to comparing the top 20 putative driver genes. Since the erythrocyte data does not come with spliced and unspliced counts, analysis with spliceJAC was not possible and CellRank had to be run in a non-default mode that does not rely on RNA velocity.

Figure 6A and B show the overlaps between the top 20 putative driver genes computed by the different methods on the two datasets. Overlaps between the two versions of of SwitchTFI are large on both datasets, indicating that the choice of the node centrality score has a limited effect on the results. Among the other methods, DrivAER has the largest overlap with SwithTFI, sharing around 50% of the top 20 drivers. For CellRank and spliceJAC, overlaps with SwitchtTFI are very small or empty, indicating that SwitchTFI returns complementary results.

**Figure 6.**
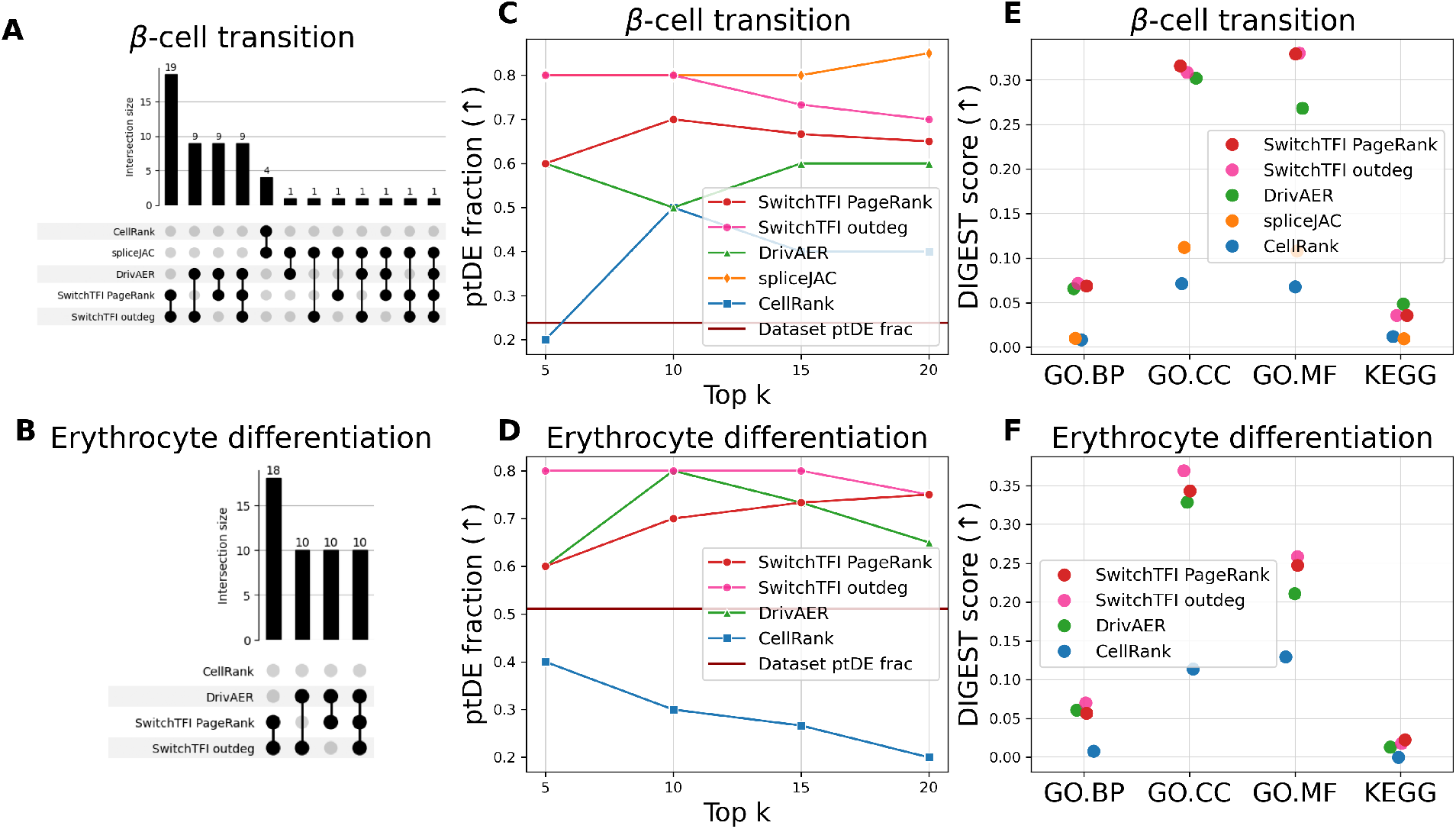
Comparison of SwitchTFI’s output to competitor methods. (A, B) UpSet plots visualizing the cardinality of the intersections of the top 20 putative driver gene sets. Empty intersections are not shown. (C, D) Fractions of ptDE genes among the top *k* = 5, 10, 15, 20 putative driver genes from each method. Higher is better. (E, F) Functional coherence scores for the sets of top 20 putative driver genes computed with DIGEST. Higher is better.

Figure 6C and D show the fractions of ptDE genes in the top *k ∈ {*5, 10, 15, 20*}* driver genes predicted by each method. On both datasets, SwitchTFI is among the best-performing methods in terms of ptDE fraction. CellRank only improves by a small margin or even fails to improve upon the fraction of ptDE genes that are in the dataset to begin with. DrivAER performs on par or slightly worse than SwitchTFI. On the *β*-cell development dataset, spliceJAC slightly outperforms SwitchTFI with respect to ptDE fraction (for the erythrocyte differentiation dataset, spliceJAC could not be used due to the lack of spliced and unspliced counts).

We further compared the list of top 20 drivers computed by the different methods with respect to functional coherence based on KEGG and GO annotations, relying on functional coherence scores which we computed with DIGEST [Adamowicz et al., 2022]. In Figure 6E and F, the functional coherence scores are compared; the full output of DIGEST can be found in Supplementary Tables 7–10 (including empirical *P*-values which show that all methods yield coherence scores which are significantly larger than expected under a random background model). Remarkably, spliceJAC achieves rather low scores here despite performing well with respect to the ptDE-based metric. In contrast, SwitchTFI and DrivAER perform well, with SwitchTFI scoring the highest. Both directly utilize information on regulatory relationships between genes to compute driver genes.

Overall, these results lead to the conclusion that all of the compared methods can predict driver genes that display potentially relevant expression trends in pseudotime. However, the functional co-herence of a proposed set of driver genes is substantially increased for methods which rely on mechanistic information, with SwitchTFI striking the best balance between ptDE fraction and functional coherence.

## 3 Discussion

We presented SwitchTFI, a GRN-based algorithm to identify transcription factors that drive cell differentiation from a progenitor to an offspring condition. We evaluated SwitchTFI on pancreatic endocrinogenesis and erythrocyte differentiation datasets and showed that it uncovers TFs known to be involved in these development processes. Moreover, SwitchTFI compares excellently against existing methods in terms of (i) functional coherence and (ii) fractions of predicted drivers that are differentially expressed over pseudotime. With respect to the latter metric, the best-performing competitor is spliceJAC, but unlike SwitchTFI, this method requires both spliced and unspliced counts as input, which are not always available. In terms of functional coherence, DrivAER is the first competitor, but is clearly outperformed by SwitchTFI when considering the fraction of differentially expressed drivers.

Despite these promising results, SwitchTFI comes with several limitations. First of all, SwitchTFI requires progenitor/offspring annotations as input. In cases where these cell state annotations do not come with the experimental procedure, they have to be inferred from the scRNA-seq data using trajectory inference tools, which may affect downstream results computed by SwitchTFI. Secondly, the quality of SwitchTFI’s results is strongly dependent on the quality of the input GRN. It is the starting point for the analysis during which edges are pruned from the GRN based on their predicted relevance to the cell state transition. Consequently, only regulatory subnetworks that are already present in the input GRN can be extracted as putative transition driver mechanisms.

For this article, we used the state-of-the-art tool Scenic to compute the baseline GRN, but inferring a high-quality from scRNA-seq is a complex problem that is not fully solved. For instance, one notable shortcoming of Scenic and other existing GRN inference methods is their stochasticity, meaning that several runs on the same data can yield different output [Ketteler and Blumenthal, 2023]. To assess to which extent this stochasticity is inherited by SwitchTFI, we conducted a small robustness study where we ran SwitchTFI multiple times on different GRNs inferred by multiple Scenic runs on the same data (see Section 4.15 for details). Indeed, differences in GRNs inferred by Scenic lead to differences in transition GRNs computed by SwitchTFI, although we observe a small increase in robustness when comparing the edges of the different networks (see Supplementary Table 11).

Future versions of SwitchTFI would hence benefit from GRN inference algorithms with improved robustness to random bias. Possible pathways toward the design of such algorithms may be to adapt SwitchTFI’s FWER control mechanism or to use network enumeration techniques as previously suggested for robust disease module mining in protein-protein interaction networks [Bernett et al., 2022]. Another interesting avenue for future work would be to develop a differential version of SwitchTFI to distinguish TFs and TF-target gene links that are relevant only to the specific transition from progenitors to offspring cells investigated by the user from those that, beyond this specific transition, are also relevant in other cell differentiation processes.

## 4 Methods

### 4.1 SwitchTFI: input data

The input to SwitchTFI consists of three components:

- A scRNA-seq count matrix *X ∈* ℝ ^*𝒸 ×ℊ*^, where 𝒸 and ℊ are sets of cells and genes, respectively.
- A GRN *G* = (*ℊ, ℰ*) that matches the biological context of the count matrix *X* or was inferred from it. *G*’s node set should match the set of genes *ℊ* in the count matrix *X* and directed edges (*i, j*) *∈ ℰ* ⊆ *ℊ ×ℊ* connect TFs *i* to their target genes *j*.
- A partition *P* ∪ *O* = 𝒸 of the set of all cells 𝒸, quantified in *X*, into progenitor cells *P* and offspring cells *O*.

### 4.2 SwitchTFI: weight fitting

Iterating over the edges *e* = (*i, j*) *∈* ℰ of the GRN *G*, a weight *w*_*e*_ *∈* [0, 1] that indicates the relevance of the respective edge to the transition between the progenitor and offspring cell states is fit to each edge as follows: The expression vectors of the TF *i* and its target *j* are the column vectors *x*_*•,i*_ *∈ ℝ* ^*𝒸*^ and *x*_*•, j*_ *∈ ℝ* ^*𝒸*^ of the expression matrix *X*. A regression tree of depth 1 (regression stump) *T*_*e*_ : *ℝ* → *ℝ* is fit to predict the target expression *x*_*•, j*_ from the TF expression *x*_*•,i*_. The fitted regression stump is defined by a decision boundary *b*_*e*_ *∈* [min(*x*_*•,i*_), max(*x*_*•,i*_)], where *b*_*e*_ is chosen such that it minimizes the mean squared error 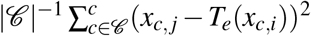. The prediction of the regression stump is defined as

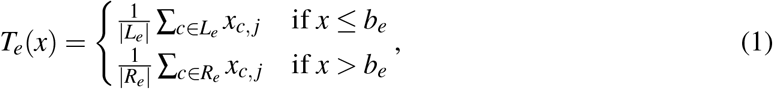

with *L*_*e*_ = *{c ∈ 𝒸* | *x*_*c,i*_ *≤ b*_*e*_*}* and *R*_*e*_ = *{c ∈ 𝒸* | *x*_*c,i*_ *> b*_*e*_*}*. The sets *L*_*e*_ and *R*_*e*_ correspond to a partition of the cells in the dataset which is compared to the partition *P, O* induced by the progenitor and offspring annotations. For this, the Jaccard index (defined as *JI*(*A, B*) =| *A*∩*B* | */* | *A*∪*B* | for sets *A, B*) is used to compute the edge relevance weight as follows:

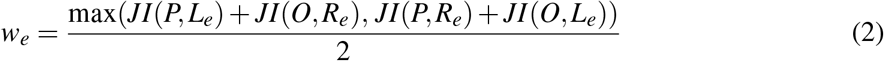

Note that the computation of *w*_*e*_ corresponds to solving a trivial 2 *×* 2 instance of a linear assignment problem, where the blocks of the two partitions are matched with respect to their Jaccard similarity. By this definition, the edge weight *w*_*e*_ is close to 1 if the partitions are similar and close to 0 if the partitions are dissimilar. Similarity of the inferred partition to the cell state partition (i.e., a high weight *w*_*e*_) is interpreted as an indicator of high relevance of the edge *e* to the transition between cell states.

Direct regulation of transcription through TFs is mainly confined to the core of individual cells. Thus, when calculating *w*_*e*_ for an edge *e* = (*i, j*), we included only cells *c* with *x*_*c,i*_, *x*_*c, j*_ *>* 0 in which both the TF *i* and its target *j* are expressed. For TF-target pairs that are both expressed only in a few cells, we observed the computed weights to behave capriciously. This problem was mitigated by including data imputation with Magic [van Dijk et al., 2018] into SwitchTFI’s preprocessing workflow. Still, we recommend not including weights *w*_*e*_ for edges *e* = (*i, j*) in the analysis where *i* and *j* are co-expressed in fewer than 20% of all cells.

### 4.3 SwitchTFI: *P*-value calculation

Only the edges most relevant to the cell state transition should be retained in the GRN. For defining a sensible cut-off for the previously computed weights, we adopted a statistical view on them. For each edge *e*, its weight *w*_*e*_ can be interpreted as the test statistic of a hypothesis test with the following null and alternative hypotheses:

- 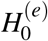: Edge *e* has no relevant explanatory power for the cell state transition from progenitor to offspring cells.
- 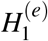: Edge *e* has relevant explanatory power for the cell state transition from progenitor to offspring cells.

The distribution and dependency structure of the weights *w*_*e*_ is unknown and we have to take into account the problem of multiple testing. Thus, we adopted the Westfall-Young permutation method [Westfall and Stanley Young, 1993] for calculating adjusted edgewise *P*-values. Again, we iterate over the edges *e* = (*i, j*) *∈* ℰ, where each edge is now associated with an already computed partition *L*_*e*_, *R*_*e*_ of the cells. Randomly permuting the progenitor/offspring labels of the cells that were used to calculate *w*_*e*_ gives a new random cell partition 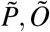 and consequently a permutation weight

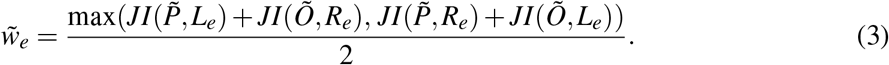

Repeating the permutation procedure *q ∈* N times results in permutation weights 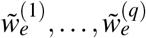 for all *e ∈* ℰ, for which we define 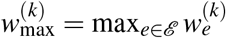 for all *k ∈ {*1,…, *q}*. The adjusted *P*-values are then computed as

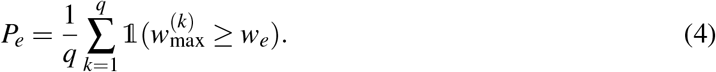

In general, if *P*_*l*_ *≤ α* for all adjusted *P*-values, this method controls the

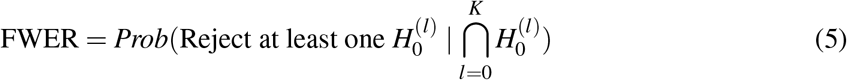

at level *α ∈* [0, 1] for *K* simultaneously performed tests with null hypotheses 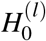 and corresponding adjusted *P*-values *P*_*l*_ [Westfall and Stanley Young, 1993]. Pruning all edges with *P*_*e*_ *> α* from *G* hence yields a cell state transition-specific GRN with controlled edge-level FWER, retaining only the most relevant edges for any kind of downstream analysis. Our recommendation is to use *α* = 0.05, as this results in a concise and well-interpretable output.

### 4.4 SwitchTFI: TF ranking

As an additional output, SwitchTFI returns a list of TFs, ranked by their relevance to the cell state transition. The ranking is computed by ordering the TFs according to their unweighted PageRank centrality [Brin and Page, 1998] in the pruned GRN. Before the PageRank algorithm is run, the direction of the edges is reversed such that they are directed from targets to TFs. This way, the TFs most central in the topology of the GRN are found by the PageRank algorithm.

Alternatively, TFs can be ranked by their weighted outdegree in the transition GRN. For this, we compute edge-level scores

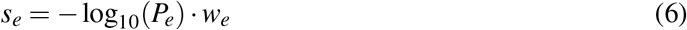

that combine the JI-based edge weights *w*_*e*_ and the Westfall-Young *P*-values *P*_*e*_ and then obtain the weighted outdegree of a TF *i* by summing up the scores *s*_*e*_ of its outgoing edges *e* = (*i, j*) in the transition GRN.

### 4.5 Datasets

The murine pancreatic endocrinogenesis dataset is part of the scRNA-seq study presented by Bastidas-Ponce et al. [2019]. It contains the transcriptomic profiles of pancreatic epithelium cells sampled at embryonic day 15.5. Cell-level annotations of developmental cell states are provided. For analysis with SwitchTFI, the dataset was subset into two separate datasets, each containing only cells from the progenitor cell state (pre-endocrine) and one offspring cell state (*α*-cell or *β*-cell).

The hematopoiesis scRNA-seq dataset was created and analyzed by Paul et al. [2015]. It contains the transcriptomic profile of myeloid progenitors differentiating towards seven distinct differentiation fates. Subsetting to the erythrocyte trajectory and merging the fine-grained cell state annotations that come with the dataset yielded the progenitor-offspring structure needed for SwitchTFI-analysis.

### 4.6 Preprocessing of scRNA-seq data

The best practices according to Ahlmann-Eltze and Huber [2023] were used as a guideline. They include filtering with respect to quality on the cell and gene level, correcting for ambient RNA, and data normalization. Additionally, we deployed the Magic [van Dijk et al., 2018] method for scRNA-seq data imputation. The individual steps are detailed and motivated in Section 2 of the supplement.

### 4.7 GRN inference

For each dataset, a GRN was inferred with the Scenic workflow presented by Aibar et al. [2017]: First, a basic GRN is inferred with GRNBoost2 [Moerman et al., 2019], which is then pruned based on *cis*-regulatory motif analysis. GRNBoost2 employs gradient boosting for more efficient GRN inference compared to the seminal GENIE3 method [Huynh-Thu et al., 2010], on which it is based. Still, the resulting GRN is coexpression-based and can contain many false positives and indirect targets. By pruning targets that lack motif support, the remaining edges constitute a biologically informed GRN. Owing to the inherent stochasticity of GRNBoost2, the output of Scenic varies greatly between individual runs. To mitigate this problem, Scenic was run *n* = 18 times and a combined GRN was constructed from the edges that appeared in *k* ≥ 9 GRNs. To perform the computations, the Python implementation pySCENIC (https://github.com/aertslab/pySCENIC) of the Scenic method was used.

### 4.8 Validation strategy

For validation steps solely concerning SwitchTFI, the input data of SwitchTFI was used to compute the validation metrics. For the method comparison, the scRNA-seq data was preprocessed according to the specifications of each method and the ptDE *q*-values were computed on non-imputed data, which otherwise had been preprocessed as previously reported. This assures a fair basis for comparison, since the competitor methods assume non-imputed input data.

The competitor methods were run as described in the tutorials that come with the respective Python packages. Also the preprocessing was adopted accordingly. The software versions and tutorials that were used are listed in Supplementary Table 1. Details on how to reproduce the results can be found at https://github.com/bionetslab/SwitchTFI-validation. Small adjustments have been made to fit our test setting. As mentioned in Section 1, for analyses with spliceJAC, the number of considered genes is limited by the minimal number of cells within a cell state. In their tutorial, only 50 genes were included. Here, 300 and 400 genes were included for the analyses of the pre-endocrine-*α* and -*β* transition. DrivAER relies on a predefined family of gene sets that is usually derived from a third-party bioinformatics database. Here, the sets of targets of individual TFs in the GRN inferred with Scenic were used to define such a family, enabling a fair comparison against SwitchTFI.

### 4.9 Pseudotime inference

The pseudotime value assigned to each cell was computed using the algorithm presented as a part of the Palantir method [Setty et al., 2019]. It is based on shortest path distances from a user-defined early cell in a graph embedding of the scRNA-seq data. The root cell was chosen according to the expression of known marker genes. For the pancreatic endocrinogenesis dataset, the cell with the highest Fev expression among the pre-endocrine cells was used; for the hematopoiesis dataset, we used the progenitor cell with the lowest Gata1 expression. The pseudotime was computed on data that was preprocessed as described before, except that the imputation step was omitted.

### 4.10 Gene expression trend computation

In accordance with Palantir [Setty et al., 2019] and CellRank [Lange et al., 2022], a generalized additive model (GAM) [Hastie and Tibshirani, 1986] with preprocessed and Magic-imputed data as an input was used to compute gene expression trends over pseudotime. In general, GAMs model the relationship between a univariate response variable, here the gene expression, and (multiple) predictor variables, here the pseudotime. Using the `LinearGAM` class of the pyGAM package [Servén et al., 2018], a GAM with a Normal error distribution and an identity link was fit. For the functional form, a spline term consisting of four basis functions of degree two was used. The prediction of the GAM for evenly spaced pseudotime values was used for visualizing gene expression trends. Moreover, 0.95-confidence intervals for the predictions as computed by pyGAM are shown around the trend.

### 4.11 Differential expression testing

Testing for differential expression of genes along a pseudotime trajectory was performed using switchde [Campbell and Yau, 2017]. A switch-like expression pattern of a gene is quantified by fitting a sigmoid function to the gene expression in pseudotime. The parameters of the sigmoid are determined as maximum likelihood estimates, where the actual optimization task is performed with L-BFGS-B [Byrd et al., 1995], a standard nonlinear optimization algorithm. A likelihood ratio test between the sigmoid model and a constant-expression model yields *P*-values for differential expression. The Benjamini-Hochberg procedure is used to compute *q*-values that are corrected for multiple testing. Furthermore, an extension to fit the sparse nature of scRNA-seq data exists, which was deployed for non-imputed data.

### 4.12 Transition GRN validation

The transition GRN *G*^*T*^ = (ℊ ^*T*^, *ℰ* ^*T*^) computed by SwitchTFI is a subnetwork of the input GRN *G* = (ℊ, *ℰ*), we write *G*^*T*^ ⊆ *G* meaning ℊ ^*T*^ ⊆ *ℊ*, ℰ ^*T*^ ⊆ *ℰ*. Indeed, SwitchTFI decides which edges *e ∈ ℰ* to retain in ℰ ^*T*^ and *G*^*T*^ is defined as the subnetwork induced by ℰ ^*T*^. Thus, random subnetworks for validation are generated by sampling |*ℰ* ^*T*^ | edges *e ∈ ℰ* uniform at random without replacement and taking the edge-induced subgraph. Repeating this procedure *q ∈* ℕ times yields subnetworks (ℊ ^(*k*)^, *ℰ* ^(*k*)^) = *G*^(*k*)^ ⊆ *G* for *k* = 1, …, *q*, which will be viewed as undirected graphs in the following.

For each of those graphs let 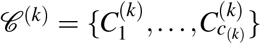 with 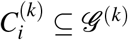 be the set of disjoint node sets that induce the *c*_(*k*)_ connected components of *G*^(*k*)^. The connectivity score of *G*^(*k*)^ is computed as

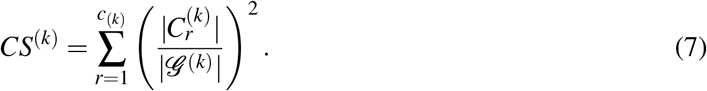

The score *CS*^*T*^ for the transition GRN is computed similarly. The corresponding empirical *P*-value is

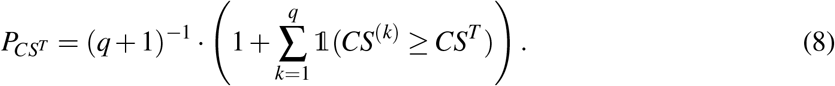

### 4.13 Gene set enrichment analysis

Gene set enrichment analysis for the top 10 predicted driver genes was performed using the Enrichr webtool [Chen et al., 2013, Kuleshov et al., 2016]. We ranked enriched terms according to their adjusted *P*-value, which is computed by Enricher with Fisher’s exact test. The enrichment was tested on the gene set libraries Gene Ontology Biological Process and Cellular Component [Ashburner et al., 2000] and Reactome [Jassal et al., 2020].

### 4.14 Functional coherence testing

Analyses of the functional and genetic coherence of the top 20 putative driver genes was performed with DIGEST [Adamowicz et al., 2022]. DIGEST takes an unordered set of genes as an input. For each gene in the set, the set of its functional annotations is retrieved from a reference database. Available databases are the Gene Ontology [Ashburner et al., 2000] (Biological Process, Cellular Component, and Molecular Function) and the Kyoto Encyclopedia of Genes and Genomes [Kanehisa et al., 2016]. The average Jaccard index for all pairs of gene-wise annotation sets is computed as the functional relevance score. To compute empirical *P*-values for the scores, randomized gene sets are created according to a random background model. Here, the fully randomized mode (genes are drawn uniformly at random without replacement) implemented by DIGEST was used.

### 4.15 Robustness study

Given an annotated scRNA-seq count matrix *X*, we computed 18 GRNs *G*_*r*_ = (𝒱_*r*_, *ℰ*_*r*_), *r* = 1, …, 18, by running Scenic 18 times on *X*. Then we ran SwitchTFI with PageRank- and outdegree-based TF ranking on these GRNs, leading to 18 transition GRNs 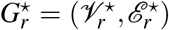 and 18 sets *T*_*r*_ of top 10 putative driver TFs in 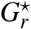. In the following, *T*_*r*_ is interpreted as an unordered set. For each type of set 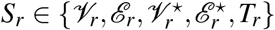, similarities *s*(*S*_1_, …, *S*_18_) are computed as the average pairwise Jaccard index

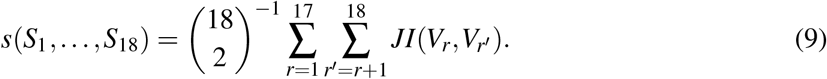

Comparing the similarity values obtained for the input GRNs *G*_*r*_ and for the transition GRNs 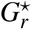 allows to assess to which extent Scenic’s lack of robustness with respect to random bias is inherited by SwitchTFI.

## Data availability

The murine pancreatic endocrinogenesis scRNA-seq dataset can be downloaded from the scVelo Python package [Bergen et al., 2020] with the `scvelo.datasets.pancreas()` function. The hematopoiesis scRNA-seq dataset can be downloaded from the Scanpy Python package [Wolf et al., 2018] with the `scanpy.datasets.paul15()` function.

## Code availability

The SwitchTFI method is available as a Python package at https://github.com/bionetslab/SwitchTFI. All other code regarding the preprocessing of the scRNA-seq data, the GRN inference, the validation, and the comparison to other methods is available at https://github.com/bionetslab/SwitchTFI-validation. Information on the software that was used is given there as well. Also, documentation and instructions on how to use SwitchTFI and reproduce results are provided.

## Author contributions statement

P.M. and D.B.B. designed SwitchTFI and drafted the manuscript. P.M. implemented SwitchTFI and carried out all analyses. A.H. and D.Z. provided critical feedback and revised the manuscript. D.B.B. supervised this work.

## Acknowledgments

P.M., A.H., and D.B.B. were supported by the German Federal Ministry of Education and Research (BMBF, 031L0309A). D.Z. was supported by the German Federal Ministry of Education and Research (BMBF, 031L0309C). Figure 1 was created with BioRender.com.

