## Supplement for "Identifying transcription factors driving cell differentiation"

Supplementary information

#### 1 In-depth review of competitor methods

**Palantir.** Palantir [Setty et al., 2019] models cell differentiation as a Markov process on a nearest-neighbor graph embedding of the scRNA-seq data. Starting from a user-defined initial cell a shortest-path based pseudotime is computed, which is used to determine the directionality of the edges in the graph embedding. Trajectories are identified by associating them with terminal states of the Markov chain. For each cell, the probability that a Markov process starting from that cell ends up in a specific terminal state is called its branch probability. The probability distribution over the trajectorial branches then serves as a soft assignment of each cell to the identified branches. For identifying key genes that drive differentiation along a certain branch towards its associated terminal state, Palantir correlates branch probabilities and gene expression using the Pearson correlation coefficient.

**CellRank.** CellRank [Lange et al., 2022] improves upon Palantir by integrating RNA velocity [La Manno et al., 2018, Bergen et al., 2020] information for inferring the directionality of the edges in the graph embedding, rendering the need for a user-defined initial cell superfluous. Again, driver genes are computed using the Pearson correlation between branch probabilities and gene expression. Both Palantir and CellRank have been proven to work well for gaining a functional understanding of cell differentiation from scRNA-seq data, but do not incorporate mechanistic information to identify genes which drive the cell differentiation.

**DrivAER.** Given a scRNA-seq count matrix, a family of gene sets, and a phenotype vector of interest DrivAER [Simon et al., 2020] combines a deep count autoencoder network (DCA) as proposed by Eraslan et al. [2019] for denoising and imputation of scRNA-seq data with a random forest (RF) model to compute relevance scores for the individual gene sets. The DCA is trained on each of the gene sets separately, such that the output of the bottleneck layer yields an embedding of the cells in low-dimensional latent space for each gene set. With these low-dimensional representations as the input and the phenotype vector as the target, a RF classification or regression model is deployed, depending on the scale of the target vector entries. The out-of-bag accuracy score serves as the relevance score for each gene set. By defining annotations of consecutive cell stages during differentiation or a pseudotemporal ordering as the phenotype of interest, the relevance score can reveal gene sets that are key to the underlying cellular process. TFs that drive differentiation can be found by defining the family of gene sets as a collection of sets of target genes of known TFs. DrivAER works well for identifying relevant TFs but cannot recover which individual TF-target gene links are relevant. This is because the embedding on whose basis the relevance scores are computed has a lower dimension (2D per default) than the number of target genes it is computed from. Individual TF-target gene relationships are lost.

**SpliceJAC.** SpliceJAC [Bocci et al., 2022] assumes a scRNA-seq count matrix with spliced and unspliced counts and a cell annotation vector containing the respective cell states as an input to infer cell state-specific gene-gene regulatory interactions and putative driver genes, using a dynamical system model. The interconnection between genes and mRNA splicing dynamics is modeled with a system of nonlinear first-order ordinary differential equations. Non-linearity appears in the gene regulation function, which represents the regulatory effects between genes with the spliced counts as the function argument. By assuming the distinct cell states to be attractors in the phase space of the underlying dynamical system and by assuming the gene regulation function to be linear when close enough to an attractor, the gene regulation function is considered to be linear for each distinct cell state. Further, by assuming the steady states of the dynamical system are fixed points of the corresponding system of differential equations, finding a solution reduces to solving a linear regression problem for each cell state. For the solution to be unique, the number of considered genes must be limited by the number of cells within a given cell state. This poses a severe limitation, since an extensive preselection of the considered genes is required. In the tutorial published with the spliceJAC Python package, only 50 out of 27998 possible genes are used for GRN inference and transition driver gene identification. Genes that are critical to the transition between cell states are found by analyzing the eigenspace of the Jacobian matrix associated with the differential equations of the starting cell state. The path displacement vector, which connects the starting and final cell state in gene space, is decomposed into a linear combination of the eigenvectors with the top $k$ largest corresponding eigenvalues. Those are regarded as unstable transition directions, such that the sum of a genes loadings in the eigenvector projection onto the transition path can be interpreted as its instability score. Genes with large instability score are thus identified as transition driver genes. Since spliceJAC focuses on individual genes, TF-target gene relations that drive differentiation cannot be identified. To mitigate this limitation, spliceJAC also provides a GRN comprised of marker genes for the initial cell state and the top transition driver genes as the vertices and the predicted interactions between them as edges. Yet, it is unclear how sensible cutoffs for the vertices and edges to be included in the GRN should be defined. Further, the edges in the GRN are inferred without utilizing transition-specific information and are thus not necessarily relevant to the cell state transition.

**Versions, access dates, and tutorials used for this study.** The versions and access dates of the competitor methods' used for this study are listed in Supplementary Table 1, along with links to the respective tutorials.

**Supplementary Table 1.** Competitor Methods, all accessed on June 14, 2024

| | Version | GitHub |
|---|---|---|
| CellRank | 2.0.4 | `https://github.com/theislab/cellrank` |
| spliceJAC | 0.0.1 | `https://github.com/federicobocci/spliceJAC` |
| DrivAER | 0.0.2 | `https://github.com/lkmklsmn/DrivAER` |

| | Tutorial |
|---|---|
| CellRank | `https://cellrank.readthedocs.io/en/latest/notebooks/tutorials/estimators/700_fate_probabilities.html` |
| spliceJAC | `https://splicejac.readthedocs.io/en/latest/notebooks/Transitions.html` |
| DrivAER | `https://colab.research.google.com/drive/1zrQ7l3Orz7h-eGEX7MHRIBTTXzL_vu9O\#scrollTo=VzAzfdHZrOWz` |

### 2 Details on scRNA-seq data preprocessing

The best practices for scRNA-seq data preprocessing according to Heumos et al. [2023] were adopted as follows:

**Quality control.** Firstly, low-quality cells were filtered. The criteria for filtering are: (1) A low number of detected genes per cell. (2) A low number of counts per cell. (3) A disproportionally high percentage of the total counts concentrated on a few, say $\leq 20$, genes. (4) A high fraction of counts from mitochondrial genes. A cell was filtered if it deviates by more than 5 median absolute deviations (mad) from the median of all cells with respect to criteria (1), (2) and (3) or more than 3 mads with respect to (4). Jointly considering criteria (1-4) reduces the risk of confounding a genuine cellular signal with low quality. The mean absolute deviation of a data vector $x \in \mathbb{R}^n$ is defined as mad $= \underset{i=1,\dots,n}{\text{median}}(x_i - \text{median}(x))$. Also cells with more than 8% mitochondrial counts were filtered. Secondly, correction for ambient RNA was performed with the SoupX [Young and Behjati, 2020] method. Lastly, quality control was also performed in the gene dimension. Genes that do not appear in at least 10 cells are considered uninformative and are filtered.

**Normalization.** The gene counts were normalized so that each cell in the dataset has the same count. To stabilize the variance across genes, the simple but effective shifted logarithm transformation $f(x) = \ln(x + 1), x \geq 0$ was applied to the normalized count matrix. Out of the 22 transformations compared by Ahlmann-Eltze and Huber [2023] the shifted logarithm performed favorably.

**Imputation.** Dropout is a common problem with scRNA-seq technologies, where counts for some genes are severely underestimated or not detected at all due to technical limitations of the laboratory procedure. The Magic [van Dijk et al., 2018] imputation method was validated to be effective at recovering gene-gene relationships [van Dijk et al., 2018, Andrews and Hemberg, 2018]. Hence it was used to preprocess the input data of SwitchTFI. Magic calculates an imputed count matrix by performing data diffusion between similar cells, the $t$th power of a distance-based cell-cell-transition probability matrix is multiplied by the count matrix. The diffusion time $t \in \mathbb{N}$ corresponds to the number of steps in the diffusion process applied to the data and thus determines the magnitude of change to the data. Here $t = 1$ was used, since for $t \geq 2$ the rate of false positive non-zero values and false positive gene-gene correlations introduced into the data increases drastically [Andrews and Hemberg, 2018].

**Supplementary Table 2.** Dataset sizes (number of cells, number of genes).

| Dataset | before preprocessing | after preprocessing |
|---|---|---|
| Pre-endocrine-$\alpha$ | 1073, 27998 | 904, 10944 |
| Pre-endocrine-$\beta$ | 1183, 27998 | 1002, 11123 |
| Erythrocytes | 1262, 3451 | 1130, 3010 |

### 3 Supplementary information on hyperparameter selection

Tables 3, 4, and 5 show the sizes of the transition GRN for different choices of the FWER threshold $\alpha \in \{0.05, 0.1, 0.2, 0.5\}$. Stricter thresholds yield smaller, more concise transition GRNs, whereas more lenient thresholds permit the inclusion of more, possibly less relevant edges. Further we examined the similarity among the sets of top $k = \{1, 5, 10, 15, 20\}$ putative driver genes across $\alpha \in \{0.05, 0.1, 0.2, 0.5\}$. The average pairwise Jaccard indices of the sets are listed in Supplementary Table 6 (averaged over $\alpha$ with $k$ fixed). It can be seen that the driver TF ranking is not greatly affected by the choice of $\alpha$. This means that the same TFs are central in the topology of the transition GRN, regardless of its size. The rankings computed with PageRank centrality are more sensitive to changes of $\alpha$, while the arguably simpler outdegree centrality can be observed to yield slightly more robust results. The choice of centrality measure is ultimately dictated by the aim of the downstream analyses. The notion of centrality and its underlying assumptions should fit the subject of study. Thus, other centrality measures such as eigenvector, katz, closeness, betweenness and vote rank as well as the option to set edge weights and change the directionality of edges are available with SwitchTFI's Python implementation.

**Supplementary Table 3.** Sizes of transition GRN for $\alpha$-cell transition data for different FWER thresholds.

| FWER threshold | # vertices | # TFs | # targets | # edges |
|---|---|---|---|---|
| 1.0, (no pruning) | 2104 | 151 | 2003 | 2732 |
| 0.50 | 348 | 42 | 316 | 434 |
| 0.20 | 325 | 39 | 296 | 402 |
| 0.10 | 303 | 31 | 281 | 382 |
| 0.05 | 284 | 28 | 265 | 359 |

**Supplementary Table 4.** Sizes of transition GRN for $\beta$-cell transition data for different FWER thresholds.

| FWER threshold | # vertices | # TFs | # targets | # edges |
|---|---|---|---|---|
| 1.0, (no pruning) | 2267 | 150 | 2175 | 3016 |
| 0.50 | 596 | 31 | 575 | 710 |
| 0.20 | 548 | 28 | 529 | 650 |
| 0.10 | 531 | 28 | 512 | 626 |
| 0.05 | 531 | 28 | 512 | 626 |

**Supplementary Table 5.** Sizes of transition GRN for erythrocyte differentiation data for different FWER thresholds.

| FWER threshold | # vertices | # TFs | # targets | # edges |
|---|---|---|---|---|
| 1.0, (no pruning) | 1008 | 60 | 984 | 1961 |
| 0.50 | 410 | 35 | 389 | 550 |
| 0.20 | 381 | 30 | 363 | 506 |
| 0.10 | 337 | 27 | 322 | 443 |
| 0.05 | 335 | 27 | 320 | 436 |

**Supplementary Table 6.** Average pairwise Jaccard indices of top-$k$ TFs across FWER thresholds (rounded to 4 decimals).

| dataset | centrality measure | $k=1$ | $k=5$ | $k=10$ | $k=15$ | $k=20$ |
|---|---|---|---|---|---|---|
| $\alpha$-cell transition | PageRank | 1.0 | 0.8334 | 0.8485 | 1.0 | 0.8194 |
| | outdegree | 1.0 | 1.0 | 0.9091 | 1.0 | 1.0 |
| $\beta$-cell transition | PageRank | 1.0 | 1.0 | 0.9091 | 0.8591 | 0.8485 |
| | outdegree | 1.0 | 1.0 | 1.0 | 1.0 | 0.9365 |
| erythrocyte differentiation | PageRank | 1.0 | 0.8334 | 0.8485 | 1.0 | 0.9091 |
| | outdegree | 1.0 | 1.0 | 0.8788 | 1.0 | 0.9365 |

### 4 Supplementary results

Supplementary Figure 1 shows the results obtained when using SwitchTFI to analyze the pre-endocrine to $\alpha$-cell transition in the pancreatic endocrinogenesis data [Bastidas-Ponce et al., 2019]. The results confirm SwitchTFI's good performance observed for the pre-endocrine $\beta$-cell transition reported in the main article. Again, the fraction of ptDE edges and vertices is significantly increased in the transition GRN over the Scenic GRN (Supplementary Figure 1A, B). Also, the joint score for the number and size of connected components can again be observed to be significantly higher for the transition GRN compared to a random background model

(Supplementary Figure 1C). Gene set enrichment for the top 10 putative driver genes revealed terms related to regulation of transcription and endocrinogenesis (Supplementary Figure 1D). Moreover, SwitchTFI compares favourably to the competitor methods also on the $\alpha$-data (Supplementary Figure 1E-G).

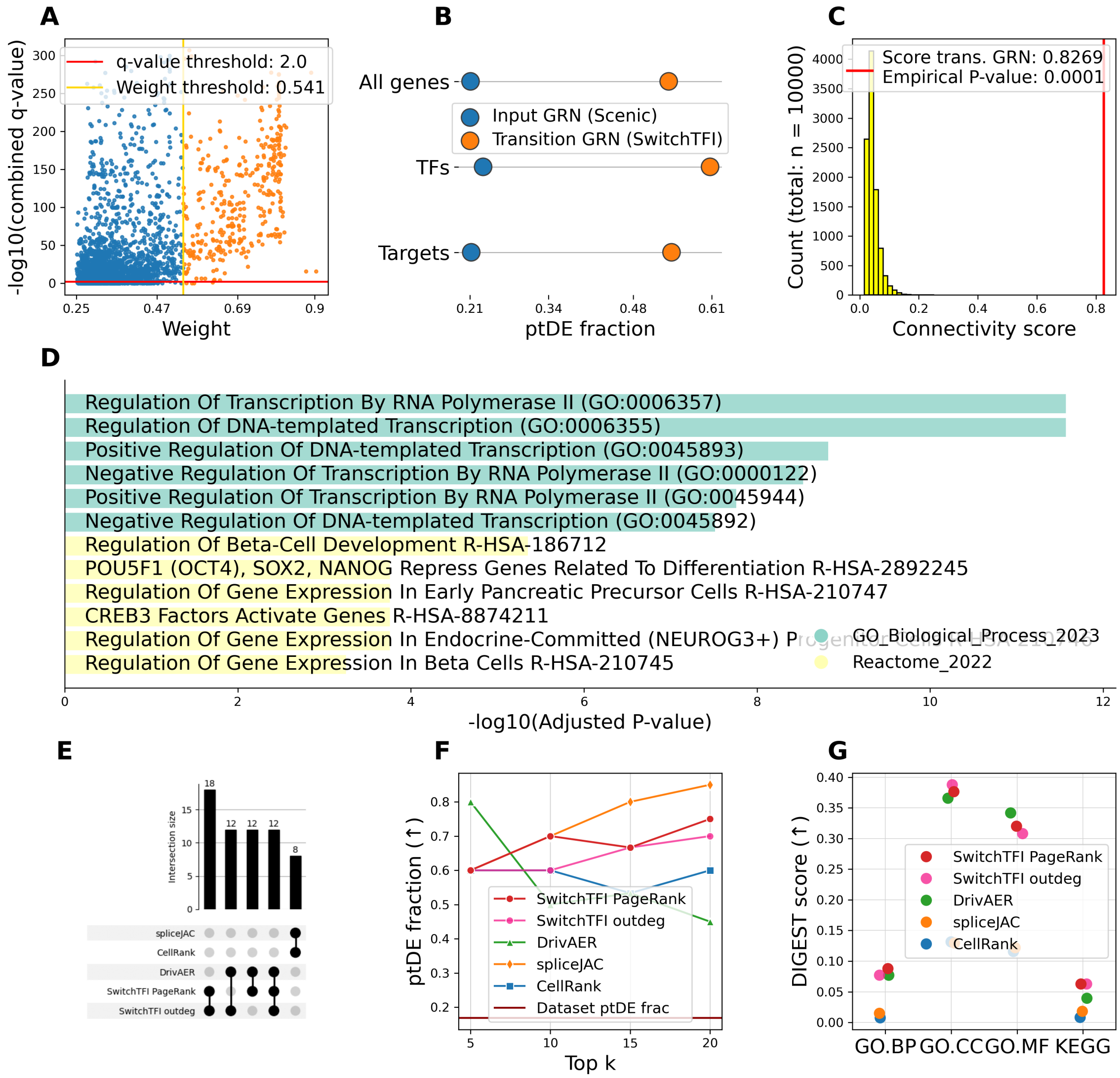


**Supplementary Figure 1.** Additional results for the pre-endocrine $\alpha$-cell transition data. (A) Edge weights vs. combined ptDE $q$-values of the unpruned Scenic GRN. Combined $q$-values are $-\log_1 0(\cdot)$-transformed. (B) Fraction of ptDE genes, TFs and targets in the Scenic and SwitchTFI GRN. (C) Histogram of the connected component scores of randomly sampled subnetworks of the Scenic GRN. The transition GRN's score is visualized as a red line. (D) Gene set enrichment results for the top 10 putative $\alpha$ driver TFs. (E-G) Method comparison results, compare Figure 5 in the main document.

Supplementary Figure 2 shows additional results for the erythrocyte differentiation dataset [Paul et al., 2015]. Here the top 10 driver genes were ranked according to their weighted outdegree in SwitchTFI's transition GRN. Figure 6 in the main document suggests that this yields slightly more relevant results for the erythrocyte differentiation data. For the top 10 driver genes predicted by SwitchTFI, their rapidly increasing or decreasing expression trends over pseudotime are displayed in Supplementary Figure 2A. For the set of top 10 driver genes (Supplementary Figure 2B), the most significantly enriched terms relate directly to hematopoiesis or more generally to the regulation of transcription (Figure Supplementary Figure 2C). For the top 10 putative driver genes, biologically validated evidence for their relevance to erythrocyte development can be found. E.g., it is well known that Gata1 and Gata2 are involved in gene regulation during erythropoiesis [Ohneda and Yamamoto, 2002]. While Gata2 is highly expressed in hematopoietic progenitors, erythroid differentiation is characterized by a switch to increased Gata1 expression [Suzuki et al., 2013]. This switch from Gata2 to Gata1 is clearly visible in the gene expression trends in Supplementary Figure 2A.

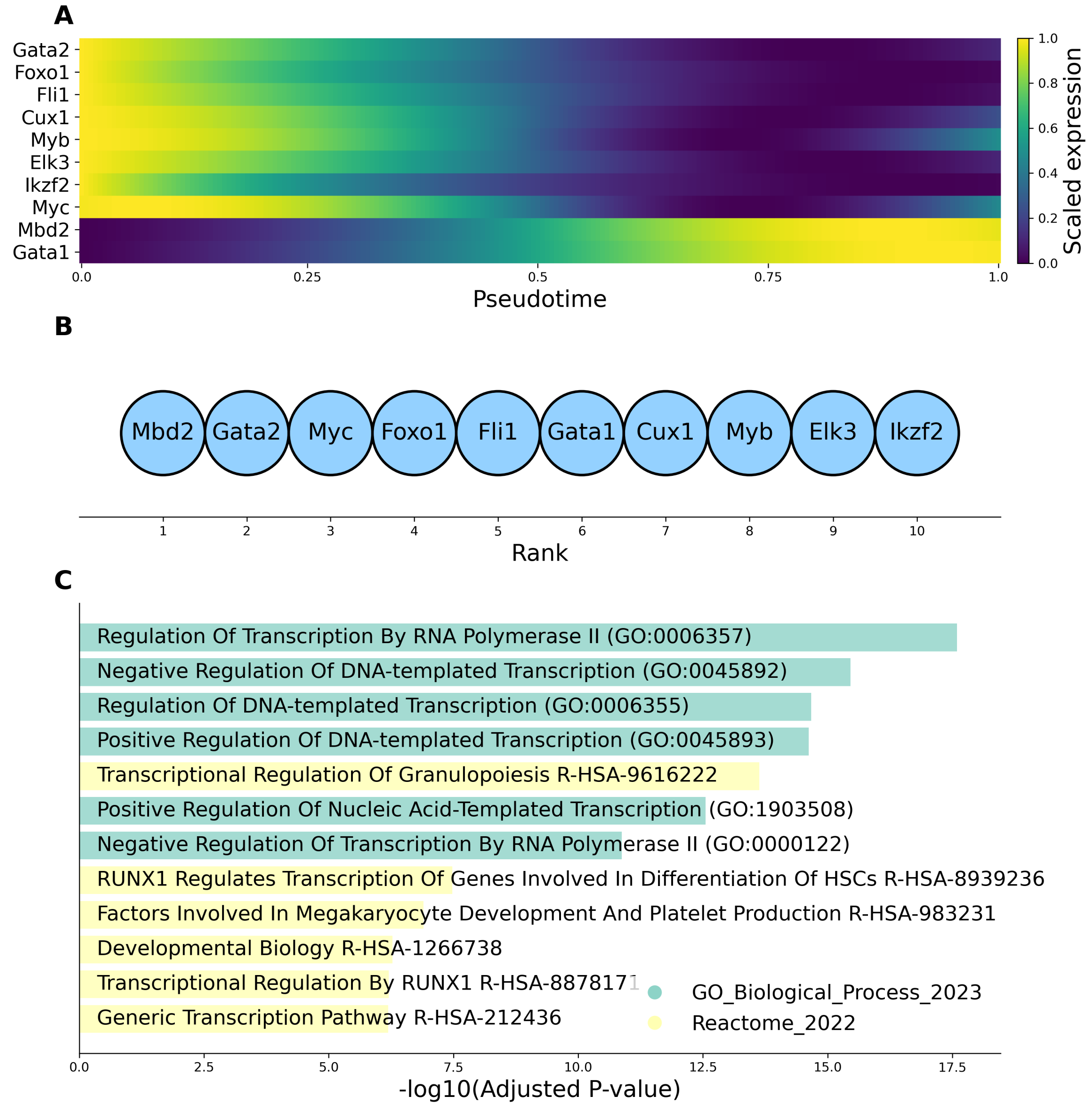


**Supplementary Figure 2.** Additional results for the erythrocyte differentiation dataset. (A) Expression trends in pseudotime of the top-10 TFs. (B) Top 10 putative driver TFs ranked by score weighted outdegree in the transition GRN. (C) Gene set enrichment results for the top 10 putative erythrocyte differentiation driver TFs.

The top 20 putative driver genes from each of the compared methods used for the method comparisons reported in Supplementary Figure 1 and in Figure 5 in the main document are listed in Supplementary Table 7. They are ordered by decreasing predicted relevance as differentiation driver genes. The detailed results of the functional coherence analysis with DIGEST are listed in Supplementary Tables 8 to 10. Lastly, the detailed results of the robustness study mentioned in the Discussion of the main document are given in Supplementary Table 11.

**Supplementary Table 7.** Putative driver genes

| Dataset | CellRank | spliceJAC | DrivAER | SwitchTFI outdeg | SwitchTFI |
|---|---|---|---|---|---|
| Pre-enocr.-$\beta$ | Ins1, Ins2, Sytl4, Ppp1r1a, Arhgap36, Nnat, Slc2a2, Ero1lb, Calr, Iapp, Gip, G6pc2, Sec24d, Pdia5, Pdia6, Hsp90b1, Papss2, Ttc28, Fkbp2, Itpkb | Pyy, Iapp, Ins2, Rbp4, Ins1, Nnat, Chgb, Malat1, Ttr, Cck, Rpl18a, Pcsk2, Actg1, Dlk1, Chga, Meis2, Runx1t1, Fev, Sec61b, Krt7 | Pdx1, Xbp1, Nkx6-1, Pax4, Ybx1, Foxa3, Fos, Etv1, Fosb, Deaf1, Pax6, Fev, Junb, Elk3, Cdx2, Nr3c1, Foxa2, Ilf2, Tbp, Rad21 | Xbp1, Pdx1, Pax4, Ybx1, Fev, Etv1, Yy1, Foxa3, Foxa2, Gata6, Jund, Lhx1, Sox4, Nr3c1, Atf3, Mnx1, Elk1, Maff, Zbtb7b, Mxd4 | Pdx1, Mnx1, Xbp1, Pax4, Ybx1, Fev, Yy1, Foxa3, Etv1, Lhx1, Nr3c1, Foxa2, Gata6, Elk1, Maff, Sox4, Jund, Atf3, Srebf2, Mxd4 |
| Pre-enocr.-$\alpha$ | Rbp4, Isl1, Pyy, Slc16a10, Peg10, Tmem27, Slc38a5, Irx2, Arg1, Lrpprc, Gast, Tmsb15l, Ank, Meis2, Anpep, Gcg, Scgn, Hap1, Ttr, Iapp | Pyy, Gcg, Iapp, Ttr, Chgb, Rbp4, Meis2, Chga, Cck, Auts2, Rps5, Tmem27, Rps9, Pcsk2, Slc38a5, Fev, Pcsk1n, Krt7, H3f3b, Mdk | Pdx1, Mnx1, Sox9, Arx, Fos, Junb, Etv1, E2f1, Pax4, Klf8, Foxa3, Clock, Jund, Taf6, Maff, Elf2, Zfp467, Foxa2, Xbp1, Fosb | Pax4, Pdx1, Creb3l2, Mnx1, Jund, Arx, Maff, Klf8, Etv1, Foxa3, Foxa2, Jun, Xbp1, Sox9, Klf3, Sox4, Ing4, Gata6, Etv5, Nkx6-1 | Pax4, Mnx1, Pdx1, Creb3l2, Jund, Cdx2, Nkx6-1, Arx, Klf8, Maff, Jun, Foxa3, Etv1, Foxa2, Xbp1, Ing4, Etv5, Sox9, Klf3, Rfx3 |
| Erythrocytes | Nusap1, Prc1, Arl6ip1, Top2a, Spire1, Mrpl47, Ckap5, Ube2c, Usp15, Rrm1, Polq, Gnl3, SMC4, Hsph1, Psmb7, Cenpf, Cenpe, Celf1, C530008M17Rik, Jmjd1c | - | Foxo1, Gata1, Mbd2, Etv6, Tfdp2, Irf1, Stat1, Sox4, Arid3a, E2f2, Irf7, Chd1, Dido1, Elk3, E2f8, Ybx1, Bclaf1, Myc, Gata2, Cebpe | Mbd2, Gata2, Myc, Foxo1, Fli1, Gata1, Cux1, Myb, Elk3, Ikzf2, Sox4, Rara, Ybx1, Irf1, Nfe2, Nfia, Tcf3, Ets1, Etv6, Tal1 | Mbd2, Foxo1, Gata2, Ybx1, Etv6, Fli1, Gata1, Myc, Cebpa, Sox4, Ikzf2, Cux1, Myb, Elk3, Taf1, Irf1, Nfe2, Pbx1, Rara, Nfia |

**Supplementary Table 8.** DIGEST results $\alpha$

| Database | Method | DIGEST score | DIGEST $p$-val |
|---|---|---|---|
| GO.BP | CellRank | 0.007578 | 0.004995 |
| | spliceJAC | 0.014933 | 0.000999 |
| | DrivAER | 0.077007 | 0.000999 |
| | SwitchTFI outdeg | 0.077127 | 0.000999 |
| | SwitchTFI | 0.087824 | 0.000999 |
| GO.CC | CellRank | 0.131471 | 0.000999 |
| | spliceJAC | 0.129550 | 0.000999 |
| | DrivAER | 0.365702 | 0.000999 |
| | SwitchTFI outdeg | 0.387706 | 0.000999 |
| | SwitchTFI | 0.376359 | 0.000999 |
| GO.MF | CellRank | 0.115743 | 0.000999 |
| | spliceJAC | 0.121951 | 0.000999 |
| | DrivAER | 0.341738 | 0.000999 |
| | SwitchTFI outdeg | 0.308004 | 0.000999 |
| | SwitchTFI | 0.320063 | 0.000999 |
| KEGG | CellRank | 0.008211 | 0.001998 |
| | spliceJAC | 0.018192 | 0.000999 |
| | DrivAER | 0.039478 | 0.000999 |
| | SwitchTFI outdeg | 0.062736 | 0.000999 |
| | SwitchTFI | 0.062736 | 0.000999 |

**Supplementary Table 9.** DIGEST results $\beta$

| Database | Method | DIGEST score | DIGEST $p$-val |
|---|---|---|---|
| GO.BP | CellRank | 0.008555 | 0.005994 |
| | spliceJAC | 0.010089 | 0.000999 |
| | DrivAER | 0.066030 | 0.000999 |
| | SwitchTFI outdeg | 0.071786 | 0.000999 |
| | SwitchTFI | 0.068847 | 0.000999 |
| GO.CC | CellRank | 0.071508 | 0.000999 |
| | spliceJAC | 0.112103 | 0.000999 |
| | DrivAER | 0.302055 | 0.000999 |
| | SwitchTFI outdeg | 0.308665 | 0.000999 |
| | SwitchTFI | 0.315826 | 0.000999 |
| GO.MF | CellRank | 0.067845 | 0.001998 |
| | spliceJAC | 0.108033 | 0.000999 |
| | DrivAER | 0.268302 | 0.000999 |
| | SwitchTFI outdeg | 0.330196 | 0.000999 |
| | SwitchTFI | 0.329223 | 0.000999 |
| KEGG | CellRank | 0.012178 | 0.002997 |
| | spliceJAC | 0.009668 | 0.000999 |
| | DrivAER | 0.048714 | 0.000999 |
| | SwitchTFI outdeg | 0.035819 | 0.000999 |
| | SwitchTFI | 0.035819 | 0.000999 |

**Supplementary Table 10.** DIGEST results erythrocytes

| Database | Method | DIGEST score | DIGEST $p$-val |
|---|---|---|---|
| GO.BP | CellRank | 0.007711 | 0.006993 |
| | DrivAER | 0.060844 | 0.000999 |
| | SwitchTFI outdeg | 0.070138 | 0.000999 |
| | SwitchTFI | 0.056724 | 0.000999 |
| GO.CC | CellRank | 0.113687 | 0.000999 |
| | DrivAER | 0.328629 | 0.000999 |
| | SwitchTFI outdeg | 0.369303 | 0.000999 |
| | SwitchTFI | 0.343111 | 0.000999 |
| GO.MF | CellRank | 0.129266 | 0.000999 |
| | DrivAER | 0.210721 | 0.000999 |
| | SwitchTFI outdeg | 0.258650 | 0.000999 |
| | SwitchTFI | 0.247286 | 0.000999 |
| KEGG | CellRank | 0.000000 | 1.000000 |
| | DrivAER | 0.012989 | 0.000999 |
| | SwitchTFI outdeg | 0.017921 | 0.000999 |
| | SwitchTFI | 0.022444 | 0.000999 |

**Supplementary Table 11.** Average pairwise Jaccard similarities for input and transition GRNs (rounded to 4 decimals).

| Dataset | vertices | | | | edges | |
|---|---|---|---|---|---|---|
| | input | trans. | top 10 TFs (PageRank) | top 10 TFs (outdeg.) | input | trans. |
| Pre-endocr. $\alpha$-cell | 0.4834 | 0.2758 | 0.5606 | 0.5507 | 0.1566 | 0.1817 |
| Pre-endocr. $\beta$-cell | 0.4956 | 0.3356 | 0.5632 | 0.5634 | 0.1607 | 0.2122 |
| Erythrocytes | 0.5991 | 0.3724 | 0.6994 | 0.7016 | 0.2553 | 0.2147 |
